# Enhanced protective efficacy of a novel, thermostable, RBD-S2 vaccine formulation against SARS-CoV-2 and its variants

**DOI:** 10.1101/2023.03.19.533338

**Authors:** Nidhi Mittal, Sahil Kumar, Raju S Rajmani, Randhir Singh, Céline Lemoine, Virginie Jakob, Sowrabha BJ, Nayana Jagannath, Madhuraj Bhat, Debajyoti Chakraborty, Suman Pandey, Aurélie Jory, Suba Soundarya S.A., Harry Kleanthous, Patrice Dubois, Rajesh P. Ringe, Raghavan Varadarajan

## Abstract

With the rapid emergence of variants of concern (VOC), the efficacy of currently licensed vaccines has reduced drastically. VOC mutations largely occur in the S1 subunit of Spike. The S2 subunit of SARS-CoV-2 is conserved and thus more likely to elicit broadly protective immune responses. However, the contribution of the S2 subunit in improving the overall efficacy of vaccines remains unclear. Therefore, we designed, characterized, and evaluated the immunogenicity and protective potential of a stabilized SARS-CoV-2 Receptor Binding Domain (RBD) fused to a stabilized S2. Designed immunogens were expressed as soluble proteins with approximately fivefold higher purified yield than the Spike ectodomain and formulated along with Squalene-in-water emulsion (SWE) adjuvant. S2 immunization failed to elicit a neutralizing immune response but significantly reduced lung viral titers in mice challenged with the heterologous Beta variant. In hamsters, SWE-formulated RS2 showed enhanced immunogenicity and efficacy relative to corresponding RBD and Spike formulations. Despite being based on the ancestral Wuhan strain of SARS-CoV-2, RS2 exhibited broad neutralization, including against Omicron variants (BA.1, BA.5 and BF.7), as well as the clade 1a WIV-1 and SARS-CoV-1 strains. RS2 sera also showed enhanced competition with both S2 directed and RBD Class 4 directed broadly neutralizing antibodies, relative to RBD and Spike elicited sera. When lyophilized, RS2 retained antigenicity and immunogenicity even after incubation at 37 °C for a month. The data collectively suggest that the RS2 immunogen is a promising modality to combat SARS-CoV-2 variants.

## INTRODUCTION

The severe acute respiratory syndrome coronavirus 2 (SARS-CoV-2) virus is the causative agent of the COVID-19 pandemic ^1^. A variety of SARS-CoV-2 vaccines were developed at an unprecedented pace to combat the pandemic ^2–8^. Spike (S) is a ∼ 200 kDa transmembrane glycoprotein composed of two functional subunits: the N-terminal S1 subunit, which mediates attachment to the host receptors, and the C-terminal S2 subunit, which facilitates membrane fusion ^9–11^. Spike protein is highly immunogenic, and the receptor-binding domain (RBD) contains the major neutralizing antibody epitopes ^12–18^. Most currently approved vaccines are based on the Spike immunogen, and a minority are based on RBD. RBD-based vaccines have been shown to elicit moderate to high-neutralizing antibody titers ^19–22^. However, due to rapid virus evolution, various variants of concern (VOC) mutations have been identified primarily in the N-terminal domain (NTD) and RBD of the S1 subunit ^23^. These mutations increase viral infectivity and induce immune evasion by evading virus neutralization. Requirements for low or ultracold temperature storage have acted as barriers to vaccine deployment in low-and middle-income countries (LMICs). There is an ongoing need for vaccination in vulnerable sections of the population, including pregnant women, those with co-morbidities and the elderly, both to protect these individuals and to minimize the potential for viral evolution in vulnerable hosts. Therefore, it remains important to develop and test vaccine candidates that confer a potent, protective, humoral immune response to a broad spectrum of SARS-CoV-2 variants and do not require a cold-chain for last mile distribution.

Compared to the S1 subunit, the S2 subunit of coronaviruses is more conserved and likely to elicit broadly protective antibodies ^24^. Several cross-reactive monoclonal antibodies against the S2 stem helix region have been identified, and a few of them have been shown to neutralize SARS-CoV-2, thereby conferring *in vivo* protection ^17,25–33^. Spike microarray analysis demonstrated that cross-reactive antibodies are elicited against the S2 stem helix region during natural infection ^17^. The stem-helix epitope in the S2 region, residues (1142-1165) is well conserved, and antibodies that bind to this region neutralize diverse coronaviruses suggesting that immunogens containing this cryptic epitope might elicit pan-coronavirus immunity.

However, the protective efficacy of S2 as an immunogen remains unclear. It is also known that the bulk of the neutralizing response is directed against the RBD. While there are also neutralizing antibodies directed against the NTD of the Spike, these have lower breadth than the RBD-directed neutralizing antibodies, as the major NTD neutralizing epitope is mutated in VOCs ^34–38^.

To address these issues, we designed a stabilized S2-ectodomain, and genetic fusions of a previously designed, stabilized RBD with S2 ^39^. These are referred to as RS2 and S2R immunogens depending on the order of connectivity of RBD and S2. Designed RS2 and S2R immunogens were expressed with ∼5.3-fold higher purified yield than stabilized Spike ectodomain in mammalian cells. Sepivac SWE^TM^ is an MF59 like oil-in-water emulsion adjuvant, subsequently referred to as SWE. Two immunizations of SWE adjuvanted RS2 induced robust neutralizing immune responses and conferred protection against SARS-CoV-2 variants in mice, showing superior immunogenicity and protective efficacy, compared to stabilized RBD, and comparable immunogenicity to stabilized Spike formulations. In hamsters, RS2 showed superior immunogenicity and protective efficacy to a similarly adjuvanted, stabilized Spike formulation. When lyophilized, RS2 retained antigenicity and immunogenicity even after incubation at 37 °C for a month. The data collectively suggest that RS2 containing vaccine formulations are a promising modality to combat SARS-CoV-2 variants.

## RESULTS

### Immunogen Design

For designing an S2-based SARS-CoV-2 immunogen, S2 subunit residues interacting with the S1 subunit were identified by accessibility calculations using the Naccess program on PDB 6VXX. Accessible surface areas (ASAs) of all residues in Spike were calculated in the absence and presence of the S1 subunit. Residues of the S2, which had a total side-chain ASA difference of >5 Å^2^, were identified as S1 interacting residues. The following mutations were made to mask newly exposed hydrophobic residues to prevent S2 aggregation in the absence of S1, namely F855S, L861E, L864D, V976D, and L984Q ^40,41^.

Proline substitutions in the loop between HR1 (heptad repeat 1) and the central helix in fusion proteins from viruses like SARS-CoV-2, SARS-CoV, MERS-CoV, and RSV are known to retain prefusion conformation, and enhance expression yields ^42–45^. More recently, HexaPro Spike, a S-6P variant, with six proline substitutions (F817P, A892P, A899P, A942P, K986P, V987P) displayed 9.8-fold increased protein expression and ∼5 °C increase in T_m_ relative to S-2P (Spike with two proline mutation) ^46^. Hence, we additionally included all these six proline substitutions in the present S2 immunogen (S2) comprising residues 698-1211 of the SARS-CoV-2 Spike. Since RBD contains the major neutralizing antibody epitopes on Spike and S2 is well conserved, we designed RS2 and S2R immunogens (**Fig. 1A**). For this, an S2 fragment comprising residues 698-1163 of the SARS-CoV-2 was genetically fused with our previously reported high-yielding, thermostable RBD containing A348P, Y365W, and P527L mutations ^20,22^.

**Fig. 1.**
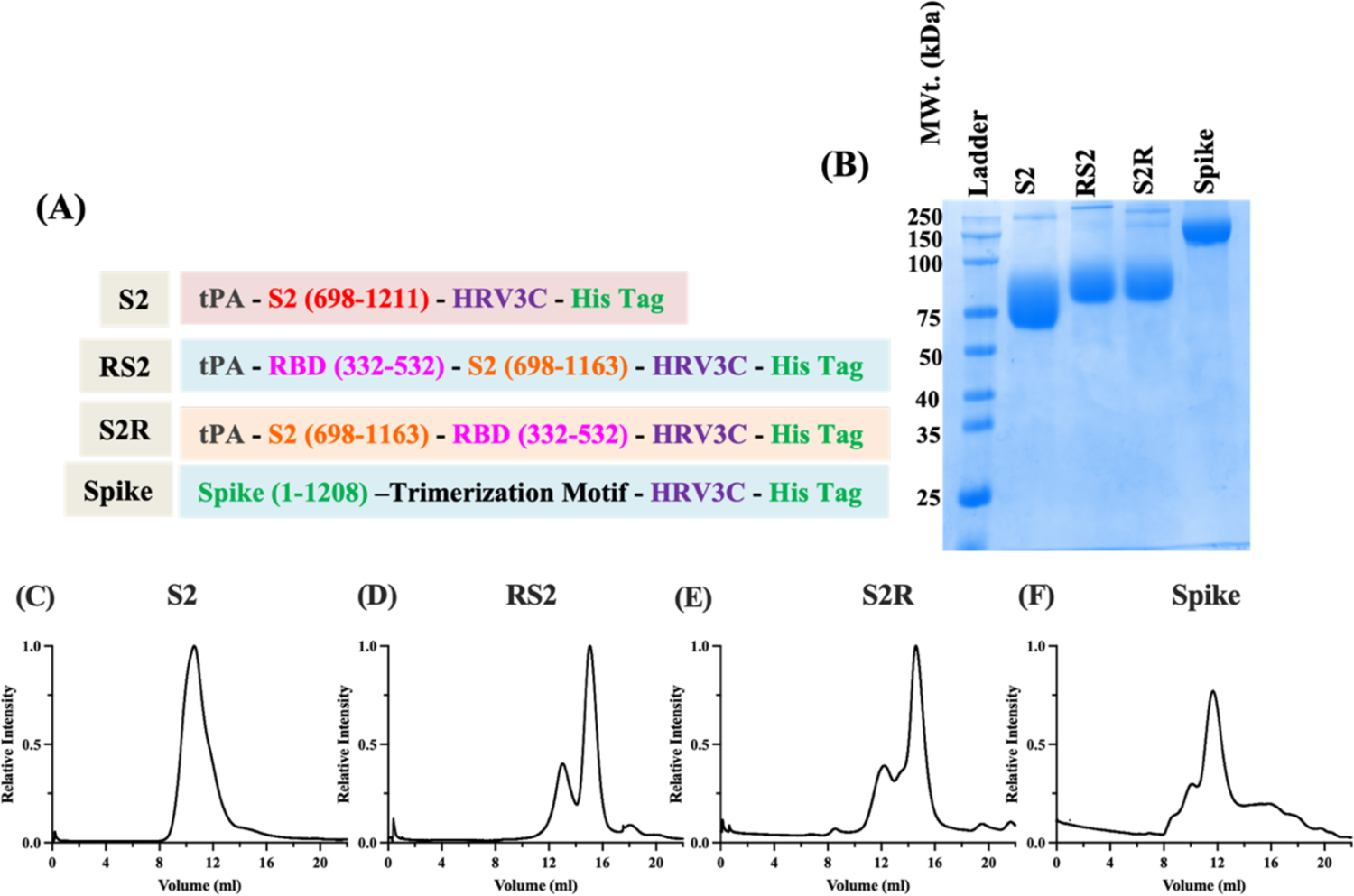
Characterization of S2, RS2, S2R, and Spike immunogens. **(A)** Schematic representation of designed immunogen sequences. **(B)** Reducing SDS-PAGE profile of protein samples. **(C-F)** SEC profile of purified **(C)** S2, **(D)** RS2, **(E)** S2R and **(F)** Spike immunogens.

### Biophysical characterization of S2, RS2, S2R and Spike

The designed immunogens were transiently expressed as secreted proteins in Expi293F suspension cells. Recombinant proteins S2, RS2, and S2R were purified in high yields of ∼120 mg/L, ∼850 mg/L, and ∼800 mg/L, respectively, using nickel affinity chromatography (**Fig. 1B**). The oligomeric state of the immunogens was determined using size exclusion chromatography (SEC), and revealed that S2 exists as homogenous nonamers in the solution (**Fig. 1C**). RS2 and S2R exist as a mixture of monomers and trimers, and the calculated molecular weights were in good agreement with the theoretical molecular weight (**Fig. 1D, E**).

For comparison, the stabilized trimeric Spike ectodomain (1-1211) containing six proline mutations (F817P, A892P, A899P, A942P, K986P, and V987P), along with the additional three RBD stabilizing mutations (A348P, Y365W, and P527L) described above, was also expressed and purified from Expi293F cells (**Fig 1A, B, F)** with a purified yield of ∼150 mg/L ^22,46^.

The apparent melting temperature (T_m_) and the thermal unfolding profile of designed immunogens and Spike was determined using Nano-DSF. The RS2 and S2R displayed similar thermal unfolding profiles and comparable T_m_ of ∼ 50 °C (**Fig. 2A and B**). The S2 immunogen exhibited T_m_ of 52.2 °C while the stabilized trimeric spike demonstrated two T_m_, T_m1_ of 50.4 °C and T_m2_ of 61.5 °C, which implied that trimeric Spike has different structural components of varying stability (**Fig. 2C and D).** RS2, S2R, and S2, when subjected to 37 °C for 1 hour, exhibited similar thermal unfolding profiles to those protein samples stored at 4°C (**Fig. 2E-G**). However, the Spike showed a slightly broadened thermal unfolding profile after incubation at 37 °C, with more noticeable broadening at 50°C (**Fig. 2H**). Moreover, RS2 and S2R were found to be stable even after incubation at 50°C, although a slight decrease was observed. On the other hand, S2 was thermally unstable at 50°C, which suggested that RS2 and S2R are more resistant to transient thermal stress than Spike and S2. (**Fig. 2E-H**). Compared to the other designed immunogens and Spike, RS2 was more rapidly digested by TPCK-trypsin at both 4 °C and 37 °C, (**Fig. 2H-K).** This data suggested that RS2 is more susceptible to proteolytic degradation than S2R, S2, and Spike, which showed highest proteolytic stability. Whether the apparent enhanced proteolytic resistance of Spike (**Fig. 2L**) is due to aggregation following initial proteolytic cleavage, remains to be elucidated.

**Fig. 2.**
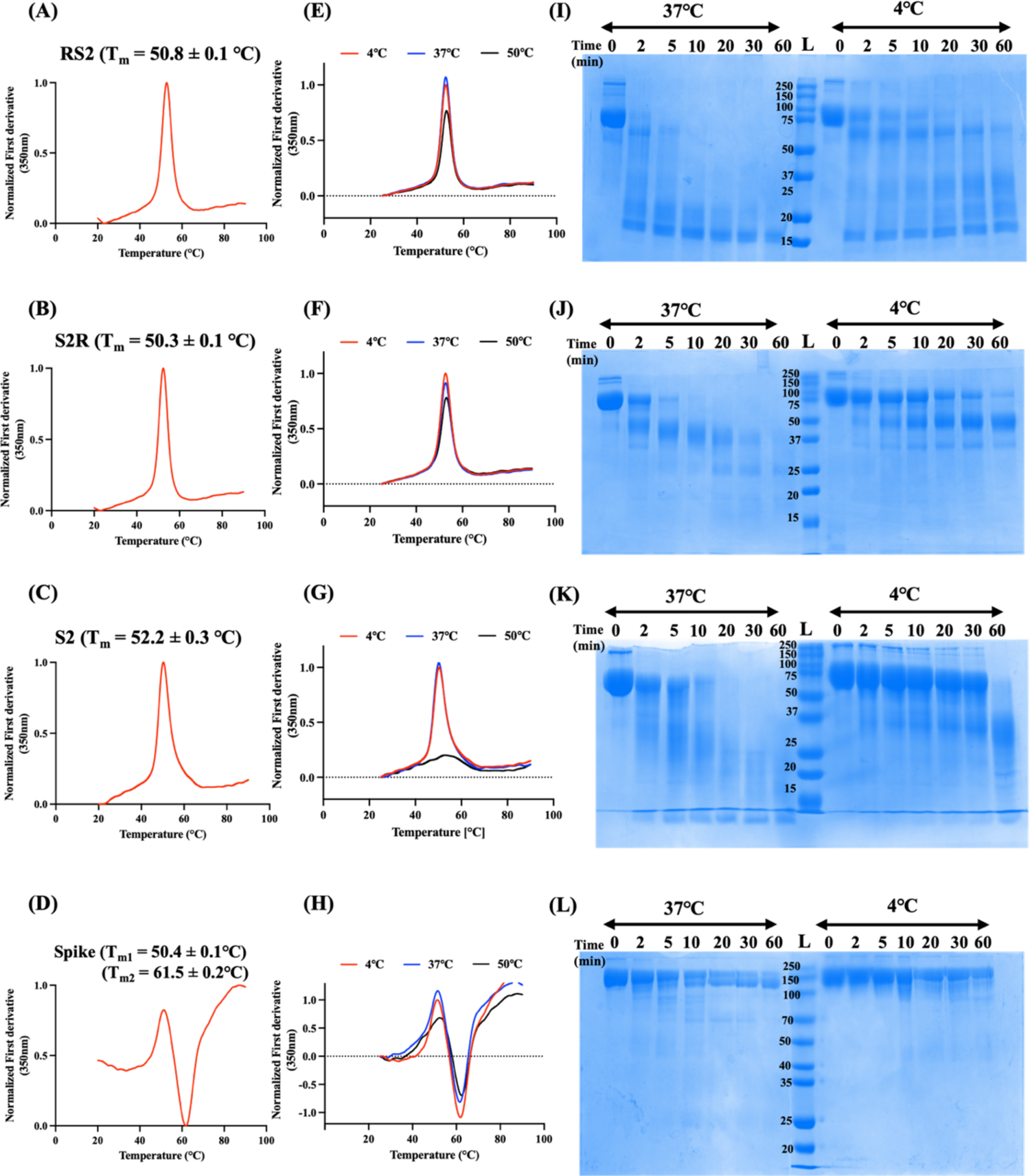
Equilibrium thermal unfolding, transient thermal stability and limited trypsin proteolytic profiles of designed immunogens. **(A-D)** Thermal unfolding profiles. Apparent melting temperature of **(A)** RS2, **(B)** S2R, **(C)** S2 and **(D)** Spike were measured using Nano-DSF. **(E-H)** Transient thermal stability profiles. Protein samples were subjected to different temperatures (4, 37, and 50 ℃) for one hour. Thermal stability of **(E)** RS2, **(F)** S2R, **(G)** S2 and **(H)** Spike was monitored using Nano-DSF. Normalized first derivative of fluorescence at 350 nm is plotted as function of temperature. **(I-L)** Proteolytic stability profile. Coomassie stained SDS-PAGE profiles of purified **(I)** RS2, **(J)** S2R, **(K)** S2 and **(L)** Spike subjected to TPCK-Trypsin proteolysis at 37 ℃ and 4 ℃.

The binding of S2, RS2, and S2R immunogens to its cognate receptor, ACE2-hFc, a panel of RBD conformation-specific (CR3022, S309, ADG-2, and H014) and S2-specific (B6 and CC40.8) antibodies were probed using surface plasmon resonance (SPR). Spike, RS2, and S2R bound well with ACE2-hFc, RBD-specific, and S2-specific antibodies (**Table 1 and 2)**. S2 binds only to S2-specific antibodies with high affinity. This indicated the proper folding of designed immunogens (**Table 2**).

**Table 1:**
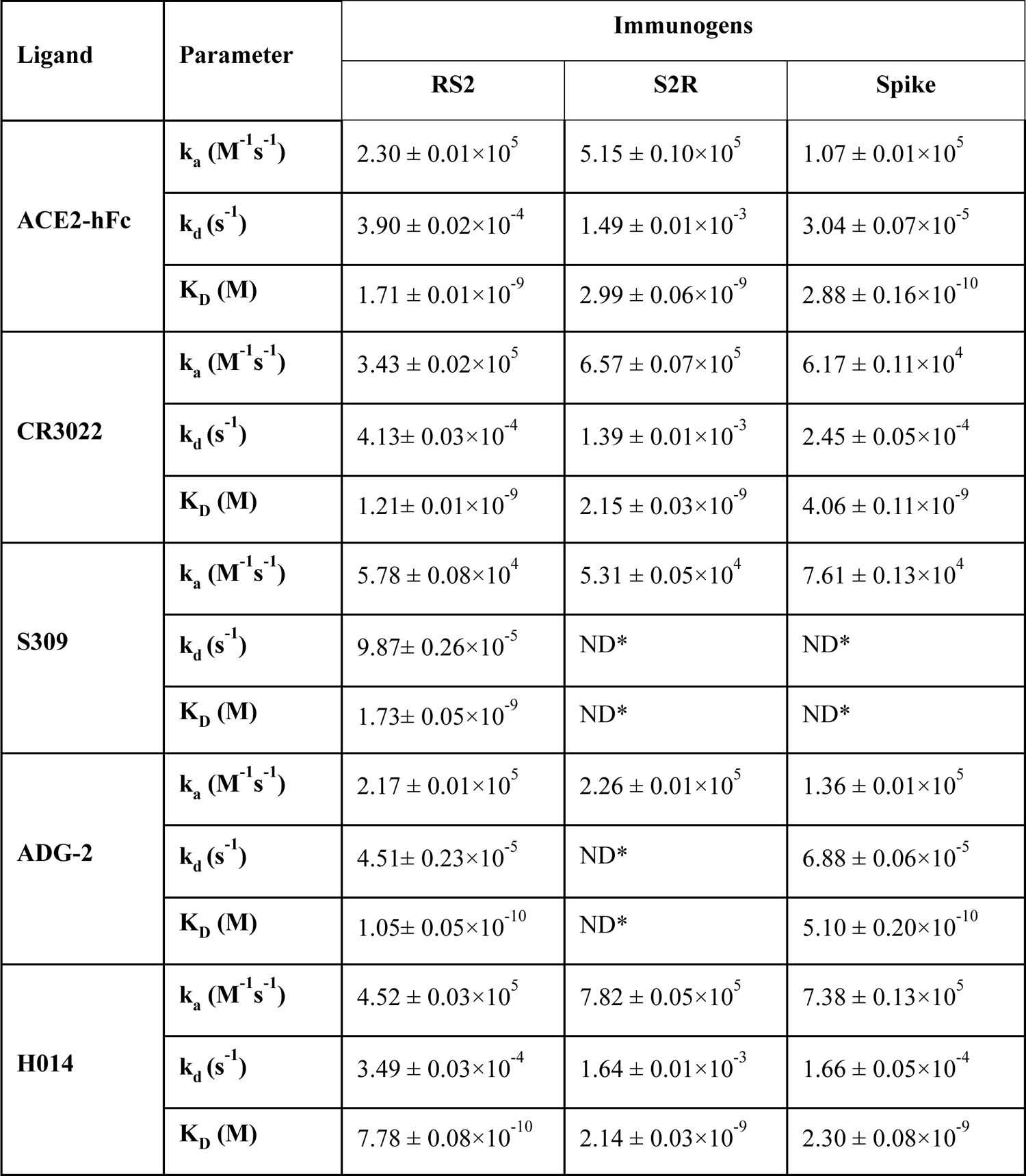
Kinetic parameters of RS2, S2R, and Spike for binding to different RBD conformation-specific ligands in PBS pH 7.4 at 25 °C. ND*: No dissociation.

| Ligand | Parameter | Immunogens |  |  |
| --- | --- | --- | --- | --- |
|  |  | RS2 | S2R | Spike |
| ACE2-hFc | $k_a$ ($M^{-1}s^{-1}$) | $2.30 \pm 0.01 \times 10^5$ | $5.15 \pm 0.10 \times 10^5$ | $1.07 \pm 0.01 \times 10^5$ |
| | $k_d$ ($s^{-1}$) | $3.90 \pm 0.02 \times 10^{-4}$ | $1.49 \pm 0.01 \times 10^{-3}$ | $3.04 \pm 0.07 \times 10^{-5}$ |
| | $K_D$ (M) | $1.71 \pm 0.01 \times 10^{-9}$ | $2.99 \pm 0.06 \times 10^{-9}$ | $2.88 \pm 0.16 \times 10^{-10}$ |
| CR3022 | $k_a$ ($M^{-1}s^{-1}$) | $3.43 \pm 0.02 \times 10^5$ | $6.57 \pm 0.07 \times 10^5$ | $6.17 \pm 0.11 \times 10^4$ |
| | $k_d$ ($s^{-1}$) | $4.13 \pm 0.03 \times 10^{-4}$ | $1.39 \pm 0.01 \times 10^{-3}$ | $2.45 \pm 0.05 \times 10^{-4}$ |
| | $K_D$ (M) | $1.21 \pm 0.01 \times 10^{-9}$ | $2.15 \pm 0.03 \times 10^{-9}$ | $4.06 \pm 0.11 \times 10^{-9}$ |
| S309 | $k_a$ ($M^{-1}s^{-1}$) | $5.78 \pm 0.08 \times 10^4$ | $5.31 \pm 0.05 \times 10^4$ | $7.61 \pm 0.13 \times 10^4$ |
| | $k_d$ ($s^{-1}$) | $9.87 \pm 0.26 \times 10^{-5}$ | ND* | ND* |
| | $K_D$ (M) | $1.73 \pm 0.05 \times 10^{-9}$ | ND* | ND* |
| ADG-2 | $k_a$ ($M^{-1}s^{-1}$) | $2.17 \pm 0.01 \times 10^5$ | $2.26 \pm 0.01 \times 10^5$ | $1.36 \pm 0.01 \times 10^5$ |
| | $k_d$ ($s^{-1}$) | $4.51 \pm 0.23 \times 10^{-5}$ | ND* | $6.88 \pm 0.06 \times 10^{-5}$ |
| | $K_D$ (M) | $1.05 \pm 0.05 \times 10^{-10}$ | ND* | $5.10 \pm 0.20 \times 10^{-10}$ |
| H014 | $k_a$ ($M^{-1}s^{-1}$) | $4.52 \pm 0.03 \times 10^5$ | $7.82 \pm 0.05 \times 10^5$ | $7.38 \pm 0.13 \times 10^5$ |
| | $k_d$ ($s^{-1}$) | $3.49 \pm 0.03 \times 10^{-4}$ | $1.64 \pm 0.01 \times 10^{-3}$ | $1.66 \pm 0.05 \times 10^{-4}$ |
| | $K_D$ (M) | $7.78 \pm 0.08 \times 10^{-10}$ | $2.14 \pm 0.03 \times 10^{-9}$ | $2.30 \pm 0.08 \times 10^{-9}$ |

**Table 2:** Kinetic parameters of S2, RS2, S2R, and Spike for binding to different S2-specific ligands in PBS pH 7.4 at 25 °C. ND*: No dissociation.

| Ligand | Parameter | Immunogens |  |  |  |
| --- | --- | --- | --- | --- | --- |
|  |  | RS2 | S2R | Spike | S2 |
| B6 | $k_a$ ($M^{-1}s^{-1}$) | $2.53 \pm 0.05 \times 10^5$ | $1.24 \pm 0.01 \times 10^5$ | $3.16 \pm 0.07 \times 10^4$ | $2.04 \pm 0.07 \times 10^5$ |
| | $k_d$ ($s^{-1}$) | $7.32 \pm 0.36 \times 10^{-4}$ | $1.53 \pm 0.8 \times 10^{-3}$ | $3.48 \pm 0.25 \times 10^{-5}$ | $2.50 \pm 0.25 \times 10^{-4}$ |
| | $K_D$ (M) | $2.30 \pm 0.12 \times 10^{-9}$ | $1.24 \pm 0.12 \times 10^{-8}$ | $1.12 \pm 0.08 \times 10^{-9}$ | $1.22 \pm 0.08 \times 10^{-9}$ |
| CC40.8 | $k_a$ ($M^{-1}s^{-1}$) | $1.62 \pm 0.01 \times 10^5$ | $1.62 \pm 0.02 \times 10^5$ | $1.19 \pm 0.02 \times 10^5$ | $9.11 \times 10^4$ |
| | $k_d$ ($s^{-1}$) | $6.77 \pm 0.01 \times 10^{-4}$ | $5.30 \pm 1.6 \times 10^{-4}$ | ND* | ND* |
| | $K_D$ (M) | $4.22 \pm 0.08 \times 10^{-9}$ | $3.23 \pm 0.03 \times 10^{-9}$ | ND* | ND* |

### Immunogenicity and protective efficacy of S2R relative to S2 and RBD immunogens against heterologous challenge

Since in the initial characterizations, S2R showed higher proteolytic stability and comparable thermal stability to RS2, the immunogenicity and protective efficacy of S2R, relative to RBD and S2 immunogens, was evaluated in hACE-2 expressing C57BL/6 transgenic mice. Our previously reported mammalian cell expressed, stabilized RBD containing A348P, Y365W, and P527L mutations was expressed and purified ^20,22^. Mice were intramuscularly immunized with 2 μg immunogens (S2R, S2 or RBD) formulated with SWE in a prime-boost regimen 3 weeks apart. SWE is equivalent to MF59, a very safe adjuvant that has been used for many years in the context of human influenza vaccines ^47^. While there are other more potent adjuvants available, stronger adjuvant mediated immune responses can be associated with unfavorable side effects ^47^. Two weeks post-boost, RBD, S2 and Spike-specific IgG titers in sera of immunized mice were measured using ELISA. Relative to RBD and S2, the S2R immunogen elicited significantly higher RBD, S2 and Spike-specific ELISA endpoint titers (**Fig. 3A-C**). S2R immunized mice elicited significantly higher neutralizing antibody titers against B.1 pseudovirus compared to RBD immunized mice, while the sera from S2 immunized mice failed to neutralize B.1 pseudovirus (**Fig. 3D**). At this administered dose, sera from RBD-immunized mice did not show neutralization against BA.1. However, S2R immunized mice sera showed neutralization against BA.1, BA.5, and BF.7 albeit at significantly reduced levels (**Fig. 3E and F**). Furthermore, these sera exhibited substantial cross-neutralization against heterologous clade1a SARS-CoV-1 viruses, which demonstrates the potential of S2R to elicit broadly protective antibodies against sarbecoviruses (**Fig. 3F**). Although the neutralization ID_50_ was too low to be measured in mice immunized with SWE formulated S2, addition of these S2 elicited sera enhanced neutralization potency of the broadly neutralizing antibody S309 (**Fig. 3G**).

**Fig 3.**
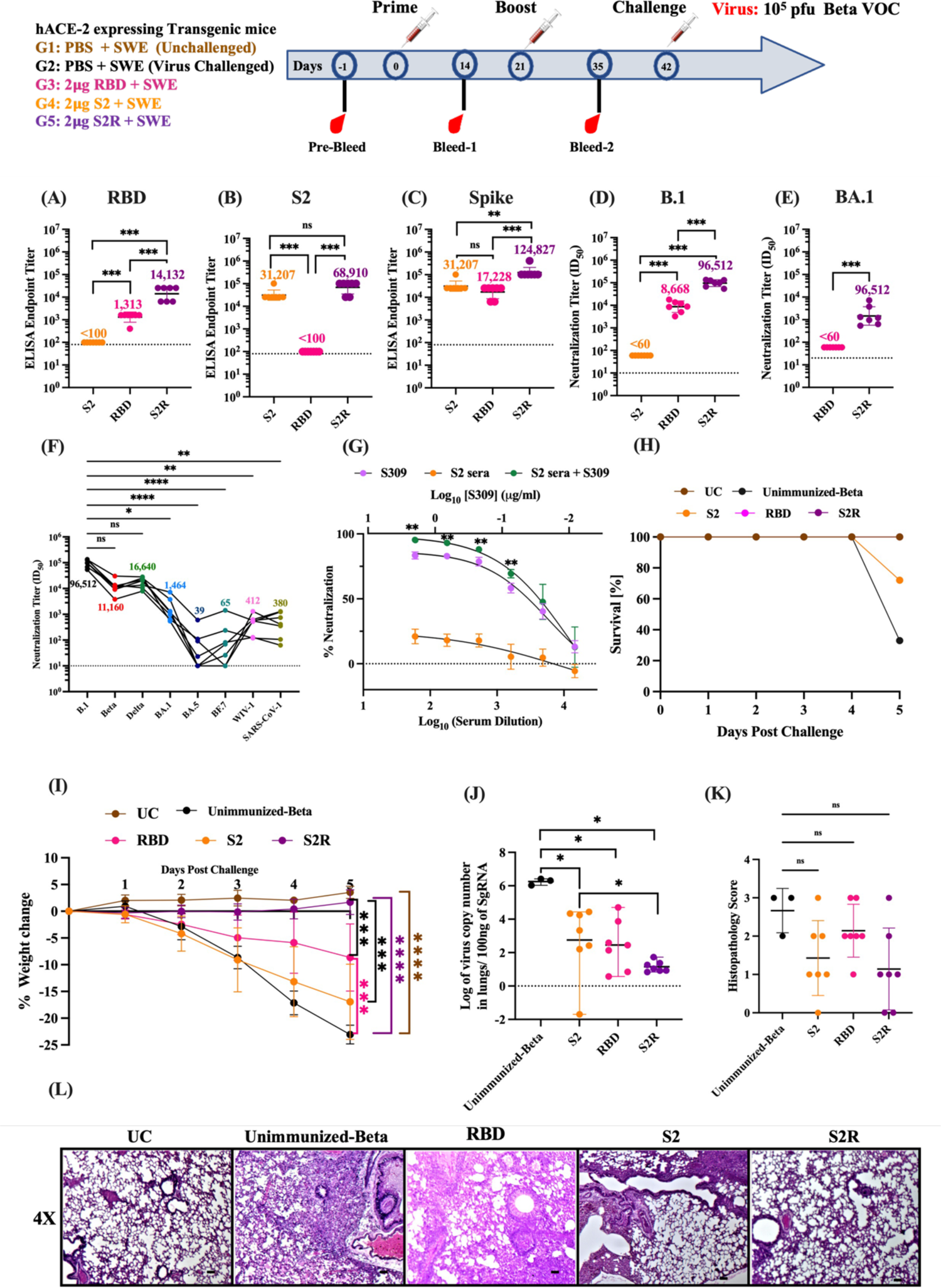
Immunogenicity of RBD, S2 and S2R in hACE-2 expressing mice. Three groups of hACE-2 expressing transgenic mice were primed and boosted with 2μg of RBD, S2 and S2R respectively, followed by an intranasal challenge with 10^5^ pfu of the beta variant of SARS-CoV-2. **(A-C)** ELISA endpoint titers against RBD, S2, and spike ectodomain respectively two weeks post-boost. **(D and E)** Neutralizing antibody titers elicited by RBD, S2, and S2R against B.1 and BA.1 Omicron SARS-CoV-2 pseudovirus. No neutralization was seen with S2 immunized animals. **(F)** Neutralizing antibody titers elicited by S2R against various pseudoviruses. Lines connect the neutralizing titers for different variants in a sera sample from an individual animal against different variants. **(G)** Neutralization curves of pooled S2 immunized mice sera, MAb S309, and S2 immunized mice sera in presence of S309. The sera sample was tested in five technical repeats. Each point represents the median of five independent values. **(H)** Survival Curve. **(I)** Average weight changes up to 5 days post-challenge. **(J)** Lung viral titer. **(K)** Histopathology scores of lungs. **(L)** Histology of lung tissue sections from unimmunized-unchallenged control (UC), unimmunized-Beta variant challenged control (Unimmunized-Beta) and mice immunized with RBD, S2 and S2R at 4X magnification. Titers are shown as geometric mean with geometric SD. The ELISA binding, neutralization titer, lung viral titer and histopathology score data were analyzed with a two-tailed Mann–Whitney test and non-parametric Kruskal-Wallis test with Dunn’s multiple correction. Pairwise weight changes were analyzed with a Multiple Student’s t-test with Bonferroni Dunn’s correction method. (ns indicates non-significant, * indicates *p* < 0.05, ** indicates *p* < 0.01, **** indicates *p* < 0.0001).

Next, the protective efficacy of RBD, S2, and S2R formulations was assessed against the Beta variant of SARS-CoV-2. Unimmunized-unchallenged, and unimmunized-Beta variant challenged mice were used as control groups. Three weeks post boost, mice were intranasally challenged with 10^5^ plaque-forming units (pfu) of Beta variant virus, and weight change was monitored for upto five days. Only 33% of unimmunized mice survived, while 72% of S2 immunized mice survived Beta-variant challenge. In contrast, all RBD and S2R immunized mice survived the Beta variant challenge (**Fig. 3H**). Post challenge, no weight change was seen in S2R immunized mice. Mice immunized with RBD, and S2 showed a significant weight reduction of ∼10 % and ∼17% respectively. As expected, no weight reduction was observed in the unimmunized group, while 22-25 % weight reduction was seen in the unimmunized-Beta variant challenged group (**Fig. 3I**). Mice immunized with either RBD, S2 or S2R showed significantly lower lung viral titers than unimmunized mice challenged with the Beta variant (**Fig. 3J**). Despite lack of neutralization, lung viral titers were significantly reduced in the S2-immunized group, suggesting that S2 provides protection by non-neutralizing mechanisms. The lung tissue sections obtained from S2R immunized mice showed clear interstitial spaces within lung epithelium and reduced immune cell infiltration compared to unimmunized-Beta variant challenged group, RBD and S2 immunized group (**Fig. 3K and L**).

### RS2 is more immunogenic than S2R in mice

At the same time, we also compared the immunogenicity of RS2 and S2R in BALB/c mice. BALB/c mice were vaccinated twice intramuscularly with either 20 μg of RS2 or S2R formulated with SWE adjuvant. Two weeks after the second vaccination, RBD and Spike specific IgG titers and neutralizing immune responses were measured in the sera samples of immunized mice. Both RS2 and S2R immunized mice elicited equivalent RBD and Spike-specific ELISA endpoint titers **(Fig. S1 A and B).** RS2 elicited significantly higher neutralizing titers against B.1 pseudovirus **(Fig. S1C)**. Neutralizing titers against Delta variant were also higher in RS2 immunized mice sera but differences did not reach statistical significance **(Fig. S1D)**. Thus, the data indicated that RS2 is more immunogenic than S2R. Hence for all further studies, including formulation stability and comparative immunization studies with Spike, RS2 was used.

### RS2 induces an equivalent neutralizing immune response to Spike ectodomain in mice and protects against mouse-adapted SARS-CoV-2 challenge

Considering that most currently licensed COVID-19 vaccines have the Spike as the sole immunogen, we compared the immunogenicity and protective efficacy of RS2 with the stabilized Spike ectodomain. BALB/c mice were intramuscularly immunized with 2 μg of RS2 or Spike formulated with SWE adjuvant in a prime-boost regimen 21 days apart. Fourteen days post-second immunization, both RS2 and Spike immunized mice elicited high RBD, S2 and Spike-specific IgG titers (**Fig. 4A-C**). Although the neutralizing titers against B.1, Beta and Delta variant pseudoviruses were comparatively higher in RS2 immunized mice sera than Spike immunized mice sera, the differences did not reach statistical significance (**Fig. 4D-F**). Sera from RS2 and Spike immunized mice also showed neutralization against BA.1, BA.5 and BF.7 pseudoviruses, albeit with lower titers (**Fig. 4G-I**). Compared to Spike, RS2 also exhibited substantial cross-neutralization against heterologous clade1a SARS-CoV-1 and WIV-1 viruses, which demonstrates the potential of RS2 to elicit broadly protective antibodies against sarbecoviruses (**Fig. 4J and K**). Three weeks post-boost, all mice were intranasally challenged with 10^5^ pfu of mouse-adapted SARS-CoV-2 MA10 virus. One-day post-challenge, mice immunized with either RS2 or spike showed a slight body weight reduction of ∼5 %, and from day two, all mice regained their initial weight. In contrast, unimmunized-MA10 challenge control mice showed ∼25 % weight loss by day four, post-challenge. No weight change was seen in the unimmunized-unchallenged control group (**Fig. 4L**). RS2 and Spike immunized mice showed significantly reduced lung viral titers compared to unimmunized-MA10 mice (**Fig. 4M**). Analysis of lung tissue sections of mice immunized with RS2 showed clear lung epithelial interstitial spaces and lower immune cell infiltration compared to both Spike immunized and unimmunized MA10 challenged group (**Fig. 4N and O**). This data suggests that RS2 elicits a broadly neutralizing humoral immune response and protects against SARS-CoV-2 MA10 virus challenge.

**Fig. 4.**
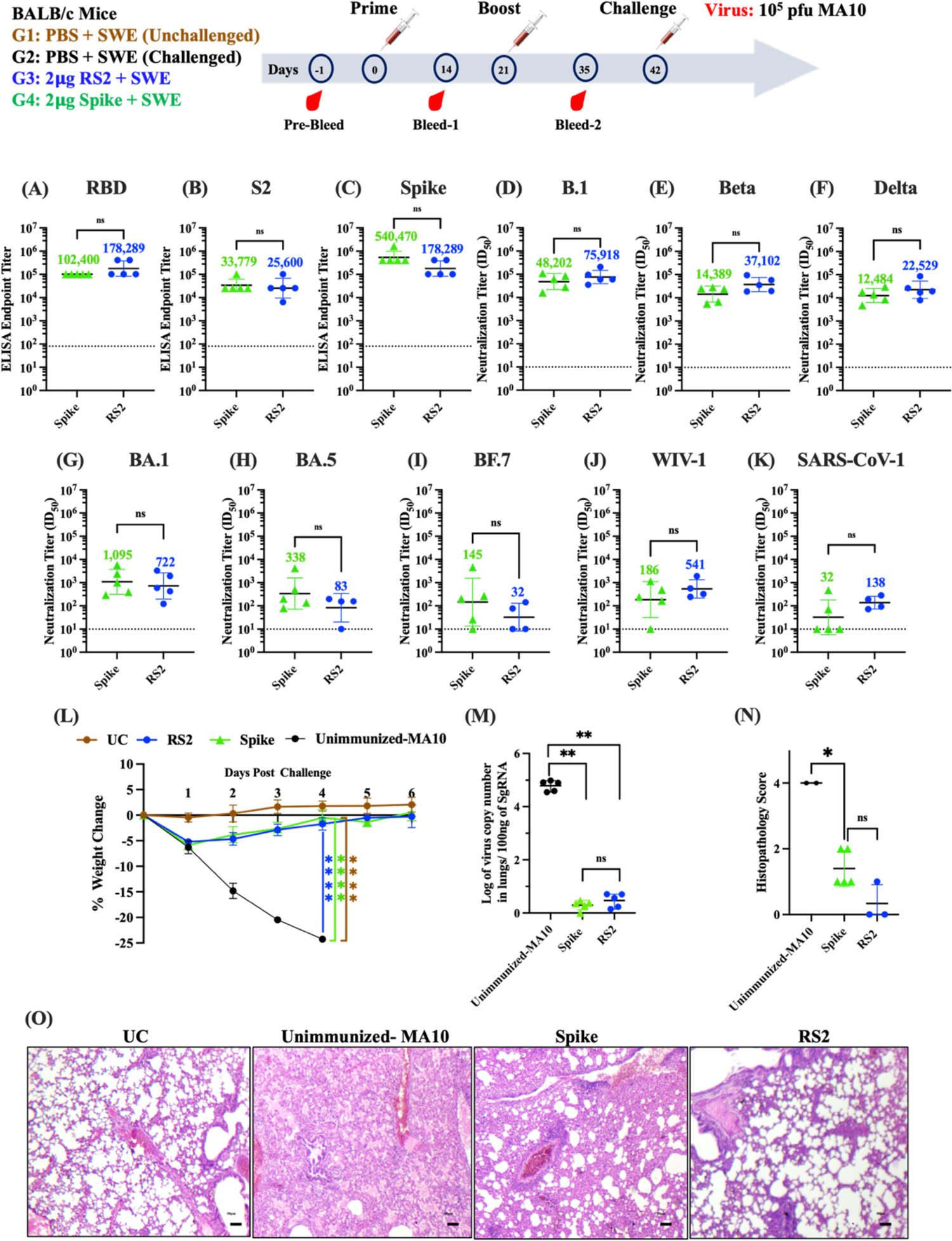
Immunogenicity of RS2 and Spike in BALB/c mice. BALB/c mice were immunized twice with either 2µg RS2 or 2µg Spike, followed by intranasal challenge with 10^5^ pfu of MA-10 mouse adapted SARS-CoV-2. Two weeks following the boost, RBD, S2 and Spike specific IgG and neutralizing titers in immunized mice sera were measured **(A-C)** ELISA endpoint titers against RBD, S2 and spike ectodomain, respectively. **(D-K)** Comparison of neutralizing antibody titers elicited by 2µg of RS2 and Spike against **(D)** B.1, **(E)** Beta, **(F)** Delta, **(G)** BA.1, **(H)** BA.5, **(I)** BF.7, **(J)** WIV-1, and **(K)** SARS-CoV-1 pseudoviruses. **(L)** Average weight changes upto six days post-MA10 challenge. **(M)** Lung viral titers **(N)** Histopathology scores of lungs. **(O)** Histology of lung tissue sections from unimmunized-unchallenged control (UC), Unimmunized-MA10 virus challenged control (Unimmunized MA-10), mice immunized with Spike or RS2 at 4X magnification. Titers are shown as geometric mean with geometric SD. The ELISA binding, neutralization titer, lung viral titer, and histopathology score data were analyzed with a two-tailed Mann–Whitney test. Pairwise weight changes were analyzed with a Multiple Student’s t-test with Bonferroni Dunn’s correction method. (ns indicates non-significant, * indicates *p* < 0.05, ** indicates *p* < 0.01, **** indicates *p* < 0.0001).

### Thermal stability of RS2

As with our previously reported RBD immunogens, our newly designed RS2 immunogen showed identical thermal unfolding profiles before and after lyophilization and solubilization, suggesting that lyophilization did not affect the thermal stability of these immunogens **(Fig. S2A)** ^20,22^. Further, the effect of transient thermal stress on the thermal stability of lyophilized RS2 was studied by performing nano-DSF. Following incubation at different temperatures (4 °C, 37 °C, 50°C, 70 °C and 90 °C) for 60 min, no change in thermal unfolding profiles was observed **(Fig. S2B)**. Moreover, the T_m_ and conformational integrity of lyophilized RS2 protein remained unchanged after a month of storage at 37 °C **(Fig. S2C) (Table 3**). In addition, the antigenicity of RS2 was assessed by performing SPR using a panel of RBD and S2-specific antibodies. Interestingly, following thermal stress, RS2 retained binding with the cognate receptor ACE2-hFc, RBD-specific antibodies (CR3022, S309), and S2-specific antibodies (B6, CC40.8). This suggests that lyophilized RS2 immunogen is resistant to transient thermal stress **(Fig. S2D-H)**. The stability of SWE adjuvanted RS2 in PBS buffer was also evaluated at 5 °C and 40 °C over a period of one month. Antigen integrity and physiochemical characterization of SWE adjuvanted RS2 formulations was carried out by performing ELISA, particle size, polydispersity, zeta potential, pH, osmolality and squalene content measurements. The RS2 formulation in SWE is stable at 5°C and 40 °C in both polypropylene tubes and glass vials for at least one month (**Fig. 5A-K**).

**Fig. 5.**
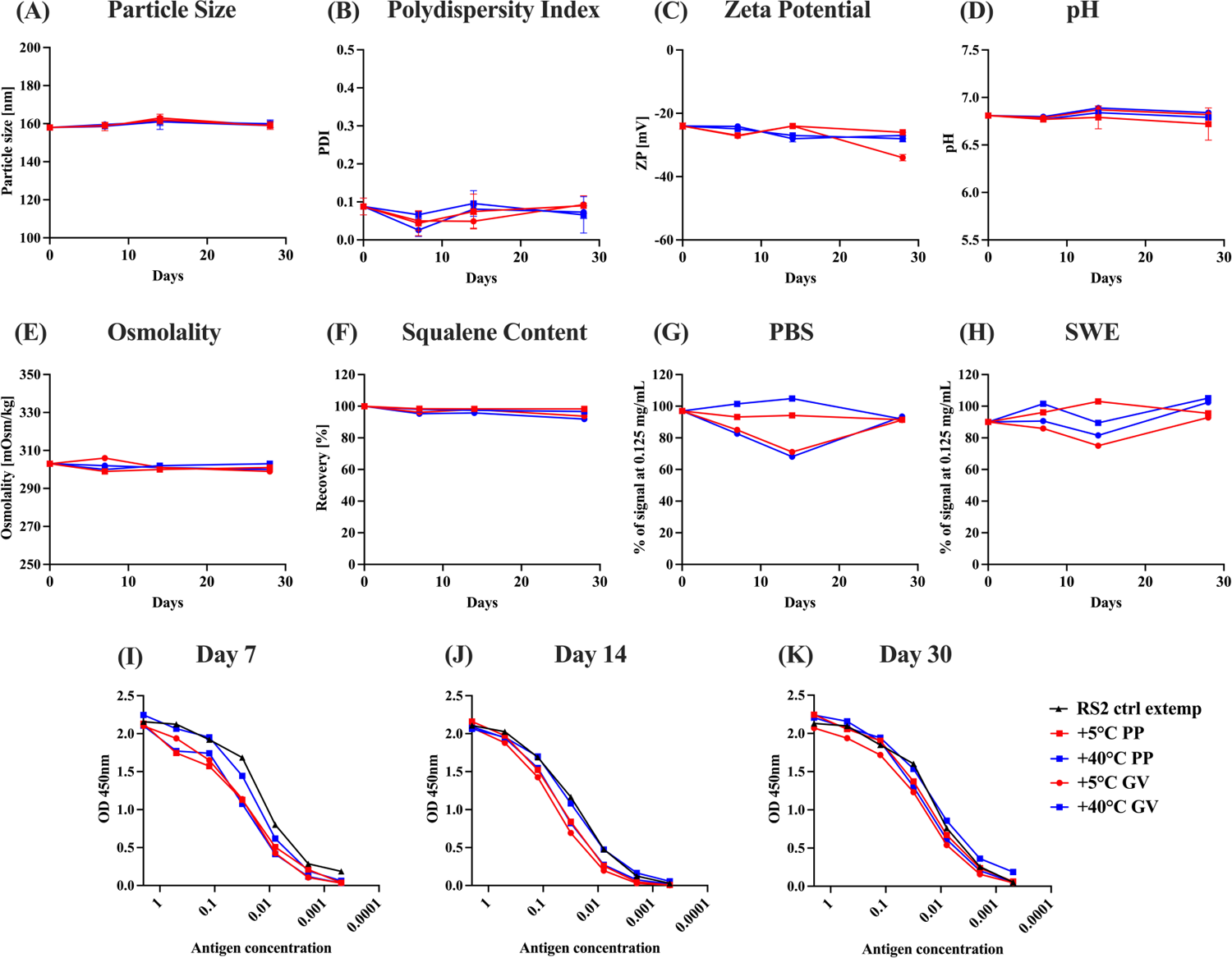
Stability of RS2. Characterization of lyophilized and resolubilized RS2 after incubation at 37 ℃ for 1 month. **(A-H)** Physiochemical characterization of RS2 in 1X PBS with equal amount (v/v) of SWE adjuvant incubated at 5 and 40 ℃ in polypropylene (PP) and glass vials (GV) for one month. Adjuvant properties were measured on day 0, day 7, day 14 and day 30. **(A)** Particle size, **(B)** Polydispersity Index, **(C)** Zeta Potential, **(D)** pH, **(E)** Osmolality, **(F)** Squalene content. **(G and H)** Antigenic integrity of RS2 in PBS and SWE adjuvant was measured based on binding to CR3022 using ELISA. **(I)** Day 7, **(J)** Day 14, **(K)** Day 30. Freshly thawed RS2 sample without any external modification ‘RS2 ctrl extemp’ was used as a control for undegraded antigen.

**Table 3:** Kinetic parameters for binding of lyophilized and resolubilized RS2 after incubation of lyophilized protein at 37 ℃ for 1 month, to different RBD-specific and S2-specific ligands in PBS pH 7.4 at 25 °C. ND*: No dissociation.

| Ligand | $k_a$ ($M^{-1}s^{-1}$) | $k_d$ ($s^{-1}$) | $K_D$ (M) |
| --- | --- | --- | --- |
| **ACE-2hFc** | $1.2 \pm 0.01 \times 10^5$ | $3.2 \pm 0.01 \times 10^{-4}$ | $2.7 \pm 0.01 \times 10^{-9}$ |
| **CR3022** | $4.8 \pm 0.01 \times 10^5$ | $7.9 \pm 0.4 \times 10^{-4}$ | $1.6 \pm 0.01 \times 10^{-9}$ |
| **S309** | $1.2 \pm 0.01 \times 10^5$ | $3.2 \pm 0.01 \times 10^{-5}$ | $2.4 \pm 0.01 \times 10^{-10}$ |
| **H014** | $5.0 \pm 0.01 \times 10^5$ | $6.8 \pm 0.4 \times 10^{-4}$ | $1.3 \pm 0.01 \times 10^{-9}$ |
| **B6** | $1.1 \pm 0.01 \times 10^5$ | $1.6 \pm 0.01 \times 10^{-5}$ | $1.2 \pm 0.01 \times 10^{-10}$ |
| **CC40.8** | $1.1 \pm 0.01 \times 10^5$ | ND* | ND* |

### Protective efficacy of lyophilized RS2 formulation after month long storage at 37 °C

The immunogenicity of the lyophilized RS2 protein stored at 37 °C for a month was evaluated in hACE-2 expressing C57BL/6 transgenic mice. Mice were intramuscularly immunized with 20 μg immunogen extemporaneously formulated with SWE adjuvant in a prime-boost regimen. Two weeks following the boost, the formulation showed high RBD, S2 and Spike-specific binding titers, and neutralizing titers against B.1, Beta, Delta, and BA.1 pseudoviruses (**Fig. 6A and B**). The above formulation elicited equivalent neutralization titers to a non-lyophilized freshly prepared RS2 formulation stored at 4 °C, against B.1 and BA.1 pseudoviruses (**Fig. 6C**). Twenty-one days following the boost, mice were challenged with 10^4^ pfu of Beta and Delta variants of SARS-CoV-2. None of the unimmunized control mice survived Beta variant challenge, while 57% of unimmunized control mice survived Delta variant challenge and the remaining mice showed 20% weight loss, nine days post-challenge. In contrast, all RS2 immunized mice survived the Beta and Delta variant SARS-CoV-2 challenges and showed no weight loss (**Fig. 6D-F**). Lung viral titers of RS2-immunized mice challenged with Beta and Delta variants were below the detection limit and lung tissues showed minimal pathology (**Fig. 6G and H).** Lung viral titers and tissue sections from the unimmunized mice challenged with the Beta variant were not examined because none of the mice survived. These findings indicate that RS2 is stable and immunogenic even after storage at 37 °C for at least one-month.

**Fig. 6.**
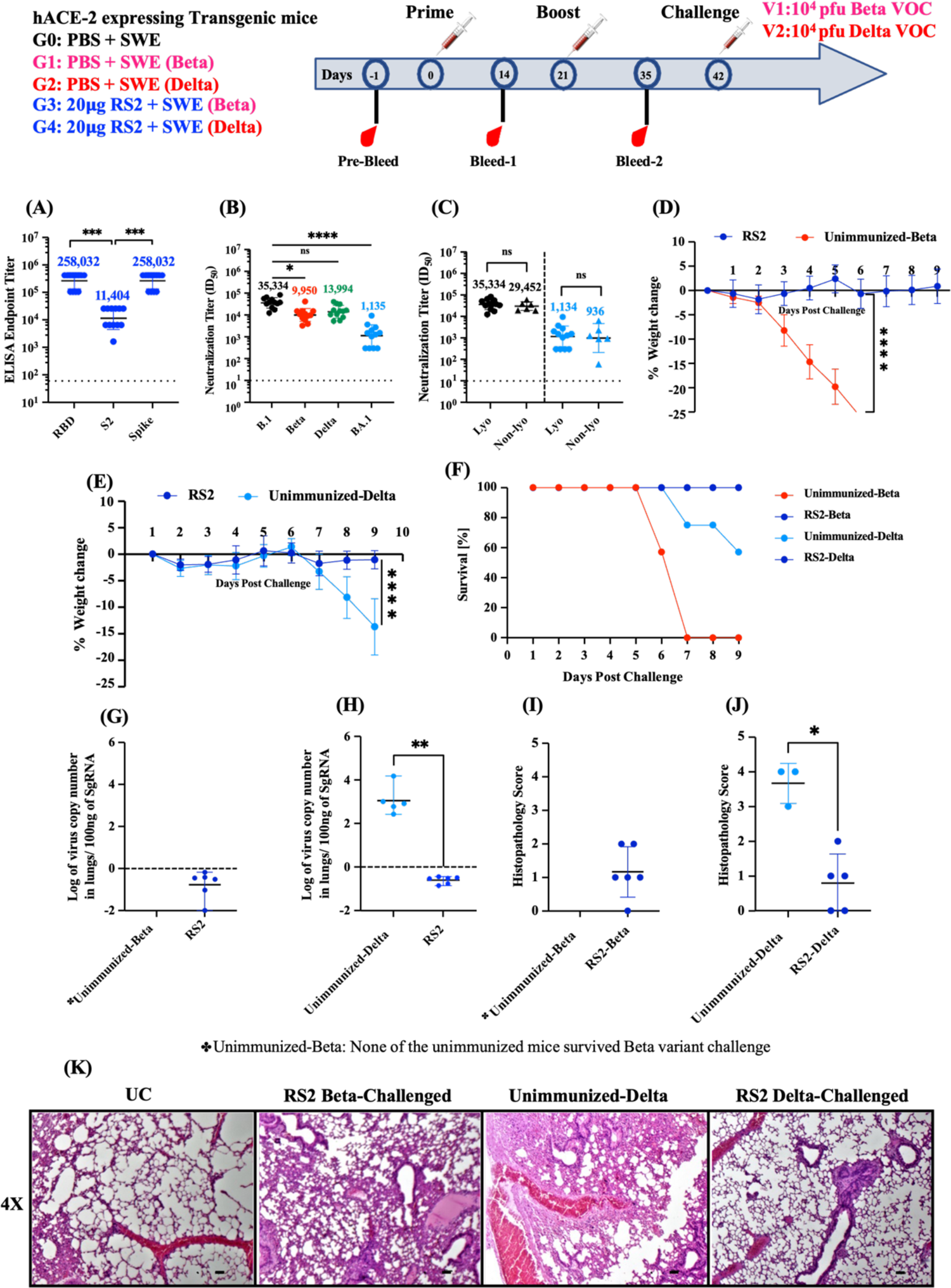
Immunogenicity of lyophilized RS2, that had been incubated at 37 ℃ for 1 month, in hACE-2 expressing transgenic mice. hACE-2 expressing transgenic mice immunized twice with 20μg of lyophilized RS2 that was previously incubated at 37℃ for over a month and then formulated in SWE adjuvant. This was followed by intranasal challenge with 10^4^ pfu Beta and Delta variants. **(A)** ELISA endpoint titers against RBD, S2, and spike ectodomain. **(B)** Neutralizing antibody titers against B.1, Beta, Delta and BA.1 pseudoviruses. **(C)** Neutralizing antibody titers elicited by lyophilized RS2 subjected to 37℃ for over a month (Lyo) and non-lyophilized RS2 (Non-lyo) stored at 4 ℃ against B.1 and BA.1 Omicron SARS-CoV-2 pseudovirus **(D and E)** Average weight change upto nine days post-Beta and Delta virus challenge respectively. **(F)** Survival curve. **(G and H)** Lung viral titers in RS2 immunized mice, challenged with Beta VOC and Delta VOC respectively. **(I and J)** Histopathology scores of lungs. **(K)** Histology of lung tissue sections from unimmunized-unchallenged control (UC), mice immunized with 20μg RS2 challenged with Beta variants (RS2-Beta challenged), unimmunized Delta virus challenged control (Unimmunized-Delta), mice immunized with 20μg RS2 and challenged with Delta variant (RS2-Delta challenged), at 4X magnification. None of the unimmunized controls survived the Beta virus challenge (Unimmunized-Beta). Titers are shown as geometric mean with geometric SD. The ELISA binding, neutralization titer, lung viral titer and histopathology score data were analyzed with a two-tailed Mann–Whitney test and non-parametric Kruskal-Wallis test with Dunn’s multiple correction. Pairwise weight changes were analyzed with a Multiple Student’s t-test with Bonferroni Dunn’s correction method. (ns indicates non-significant, * indicates *p* < 0.05, ** indicates *p* < 0.01, **** indicates *p* < 0.0001).

### RS2 shows superior immunogenicity and protective efficacy to stabilized Spike ectodomain in hamsters

Protective efficacy of RS2 was compared with the stabilized Spike against challenge with the Beta variant of SARS-CoV-2 in Syrian hamsters. Female Syrian Golden hamsters were immunized with 5μg of stabilized Spike or RS2 formulated with SWE on day 0 and day 21, while the control group of hamsters was immunized with SWE adjuvant in PBS. Both Spike and RS2 immunized hamsters elicited high RBD, S2 and Spike-specific ELISA endpoint titers (**Fig. 7A-C**). Notably, two immunizations with 5 μg SWE adjuvant formulated RS2 elicited significantly higher neutralizing titers against B.1, Beta, Delta and Omicron BA.1, BA.5 and BF.7 variants than corresponding titers elicited by SWE formulated Spike in hamsters (**Fig. 7D-I**). Consistent with the BALB/c mice study, RS2 immunized hamsters exhibited substantially higher cross-neutralizing activity against clade 1a sarbecoviruses, WIV-1 and SARS-CoV-1 compared to those of Spike immunized hamsters (**Fig. 7H and K**). Moreover, hamsters immunized with RS2 showed initial transient weight loss (up to 3 %) and regained weight at 3 days post Beta variant infection. In contrast, Spike-immunized hamsters showed comparatively higher lung viral titers and weight loss, while no weight regain was observed (**Fig. 7L and M**). Both RS2 and Spike immunized hamsters showed significantly reduced lung viral titers compared to unimmunized-Beta challenged hamsters (**Fig. 7M**). Analysis of RS2 and Spike immunized hamster lung tissue sections showed clear lung epithelial interstitial spaces and lower immune cell infiltration compared to the unimmunized Beta-challenged groups (**Fig. 7N and O**). Overall, the data shows that RS2 is more immunogenic and efficacious than Spike in hamsters.

**Fig. 7.**
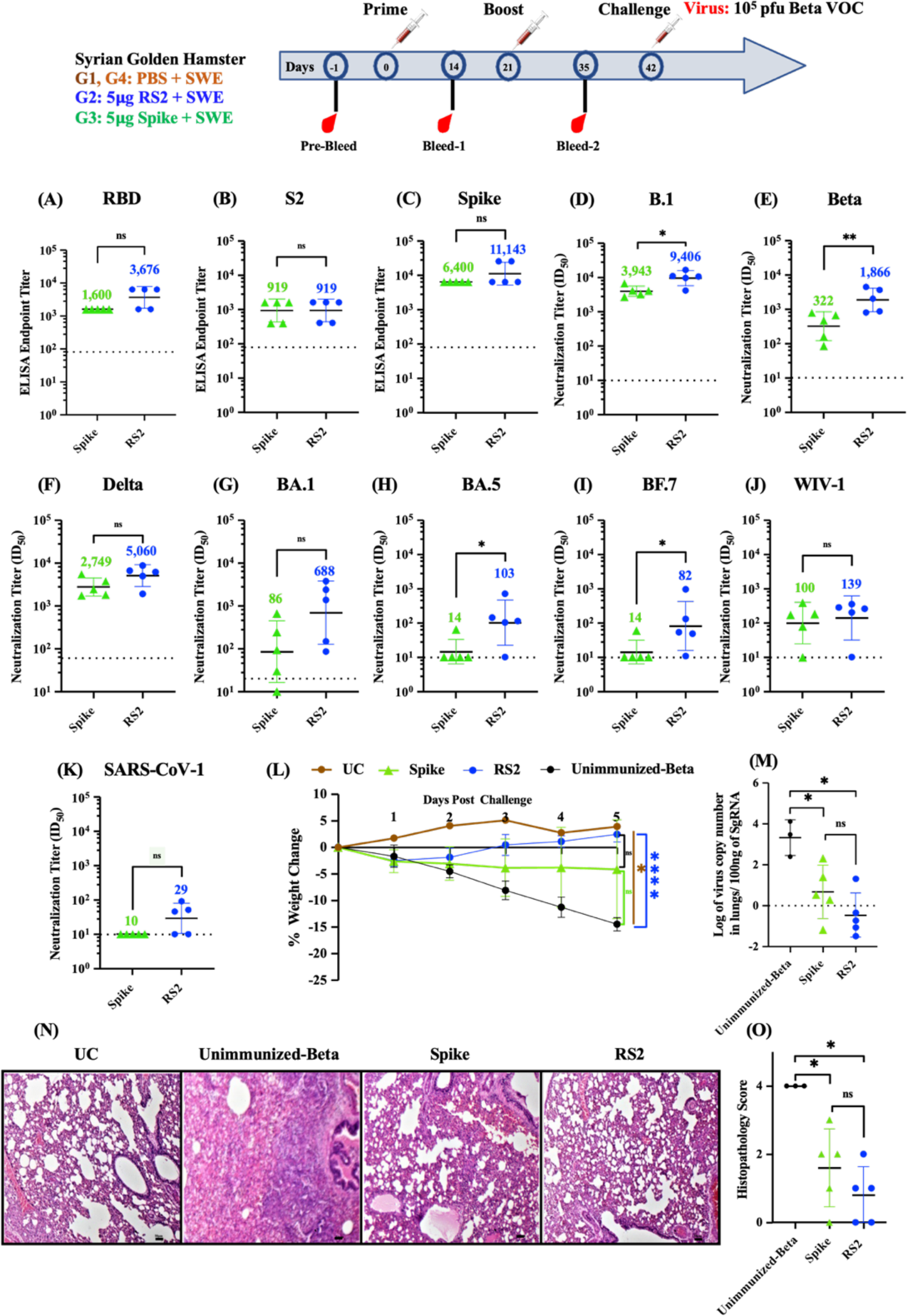
Comparative protective efficacy of RS2 and spike in hamsters. Syrian hamsters were immunized twice with 5µg of RS2 or Spike, followed by intranasal challenge with 10^5^ pfu of the Beta VOC. **(A-C)** ELISA endpoint titers against RBD, S2 and spike ectodomain, respectively. **(D-K)** Comparative neutralizing antibody titers elicited by 5µg of RS2 and spike against **(D)** B.1, **(E)** Beta, **(F)** Delta, **(G)** BA.1, **(H)** BA.5, **(I)** BF.7, **(J)** WIV-1 and, **(K)** SARS-CoV-1 pseudovirus. **(L)** Average weight change upto five days post-beta variant virus challenge. **(M)** Lung viral titers **(N)** Histology of lung tissue sections from unimmunized-unchallenged control (UC), unimmunized-Beta challenged control (Unimmunized-Beta) and hamsters immunized with Spike and RS2 at 4X magnification. **(O)** Histopathology scores of lungs. The ELISA binding, neutralization titer, lung viral titer and, histopathology score data were analyzed with a two-tailed Mann–Whitney test. Pairwise weight changes were analyzed with a Multiple Student’s t-test with Bonferroni Dunn’s correction method. (ns indicates non-significant, * indicates *p* < 0.05, ** indicates *p* < 0.01, **** indicates *p* < 0.0001).

## DISCUSSION

Vaccination has significantly reduced the global health burden caused by the COVID-19 pandemic. While vaccines have been proven to be highly efficacious against ancestral SARS-CoV-2 stain, their efficacy has rapidly declined against new VOCs and periodic updating of vaccines appears to be necessary. Most VOC mutations are identified in the receptor binding motif (RBM) region of RBD. The substantial waning of serum-neutralizing antibody titers and the emergence of variants with increased transmissibility and neutralizing antibody escape responses are associated with increased breakthrough infection in vaccinated individuals. To curb infection and sustain vaccine effectiveness, booster doses are required.

Various approaches have been deployed to broaden the protection breadth of SARS-CoV-2 vaccines, which include mosaic nanoparticle vaccine designs that display multiple RBDs from different sarbecoviruses, and conferred broad protection against diverse coronaviruses, albeit with a reduction in potency against some SARS-CoV-2 VOCs ^48–50^. While promising, such nanoparticle vaccine designs elicit titers against the nanoparticle scaffolds, and employ a large number of antigens, which adds to manufacturing complexity. Alternatively, vaccine candidates based on conserved antigens like S2 are also reported ^51–53^.

In the present study we designed a stabilized S2 ectodomain, and the genetic fusion of RBD and S2. The RS2 and S2R contain the RBD component to elicit potent neutralizing antibodies, and the S2 to increase immunogenicity and protective breadth. The novel RS2 and S2R immunogens exhibited ∼5.3-fold higher protein yields and were shown to be more stable to transient thermal stress than a stabilized Spike (**Fig. 2E and H**).

All the currently licensed COVID-19 vaccines require refrigerated or frozen storage. Recently, Ferritin displayed, alum adjuvanted Spike construct also displayed promising immunogenicity in non-human primates and could be stored at 37°C for at least two weeks^54^. In the present study we have shown that our RS2 vaccine candidate can retain its thermal stability and antigenicity without any loss of immunogenicity and protective efficacy at 37 °C for at least a month. Encouragingly, RS2 immunogen formulated with SWE was also shown to maintain its physico-chemical stability at 40°C for up to one month, however the immunogenicity needs to be evaluated. This exceptional stability of the RS2 candidate will facilitate distribution in low resource settings and will reduce the cost associated with low or ultracold temperature storage and transportation.

Despite being based on the original Wuhan strain sequence, RS2 elicited neutralizing antibodies against B.1, Beta, Delta, Omicron (BA.1, BA.5, BF.7) and clade 1a SARS-CoV-1 and WIV-1 pseudoviruses in mice and hamsters. While many SARS-CoV-2 immunogens exhibit high immunogenicity in mice, immunogenicity is poorer in hamsters, and more predictive of immunogenicity in humans^55–58^. Therefore, we chose to use the hamster animal model to further probe for apparent differences in immunogenicity between RS2 and Spike immunized animals. In hamsters, the RS2 immunogen elicited significantly higher neutralizing titers against all the tested VOCs compared to Spike (**Fig.7D-K)**. Qualitatively, RS2 immunized animals also exhibited less weight loss and lower lung viral titers compared to Spike immunized animals (**Fig.7H-J)**. However, due to the limited number of animals in the study, the differences in weight loss and lung viral titers between the two groups did not reach statistical significance. In addition, the sera obtained from mice immunized with RS2 showed competition with S2X259, and B6 antibodies. In contrast, the sera from mice immunized with Spike or RBD failed to show competition with these antibodies **(Fig. S3)**. These findings suggest that RS2 sera specifically target highly conserved epitopes located in the RBD and S2 stem helix, which are likely less accessible in Spike or RBD based immunogens. In contrast to Spike where regions of RBD that contain neutralizing epitopes are occluded in the down conformation, there no such conformational constraint in the RBD-S2 fusions.

Like other vaccine formulations, it will be necessary to periodically update formulations based on RS2. However, due to the lower mutation rate in the S2 region and higher yield, updating RS2 vaccine sequences to match circulating vaccines is expected to be easier than updating full length Spike based vaccines.

In contrast to RS2, the S2 component alone induces high IgG binding titers but sera failed to neutralize the viruses^52,53^. Addition of S2-elicited sera significantly enhanced neutralization potency of the RBD directed, broadly neutralizing S309 antibody (**Fig. 3G).** This finding further supports the role of S2 in boosting immunogenicity and enhancing the protective efficacy of the vaccine designs based on genetic fusion of RBD and S2 as in RS2 and S2R.

Similar to previous reports, our study reaffirms that despite lack of significant neutralization titers, lung viral titers were significantly reduced in S2 immunized animals. This indicates that either T-cell mediated or S2 directed antibodies, could offer protection through non-neutralizing mechanisms, but these were not evaluated in the present study. In future, further characterization of cellular immune responses and the effector functions of S2 directed antibodies will be evaluated, to provide insights into the diverse mechanisms underlying the observed reduction in viral loads.

In conclusion, our study demonstrates that RS2 has several advantages compared to Spike, based on its improved immunogenicity in small animals, higher thermal tolerance, and substantially greater purified yield. Currently a significant proportion of the world’s population has been immunized with vaccines with full-length Spike immunogens, and likely an even larger number of people have acquired natural immunity through infection. In this scenario an RS2 vaccine could still be used as a booster vaccine in vulnerable groups that will require annual vaccination and could also be combined with other vaccine formulations for respiratory viruses. Given that the virus is likely to continue to circulate in humans for the foreseeable future, it is important to continue developing and improving vaccine technologies both to combat this evolving virus and for future preparedness against other coronavirus pandemics. The RS2 vaccine design can be used as a prototype for designing vaccines that target other coronaviruses, or potential future variants of SARS-CoV-2.

## MATERIALS AND METHODS

### Expression and Purification of recombinant proteins

Designed immunogens S2, RS2, S2R and Spike are based on the ancestral Wuhan SARS-CoV-2 strain (GenBank Id: YP_009724390.1). Mammalian codon optimized genes for S2, RS2 and S2R were synthesized at GenScript Inc. The designed immunogens consisted a HRV3C protease site (LEVLFQGP) at the C-terminus of proteins to facilitate histidine tag removal after purification. Recombinant proteins S2, RS2, S2R, RBD, and Spike were transiently expressed in Expi293F™ cells according to the manufacturer’s guidelines (Gibco, ThermoFisher Scientific). Briefly, Expi293F™ cells were maintained at a cell density of ∼3 × 10^6^ viable cells/mL in the Expi293F™ expression medium. Plasmid DNA and ExpiFectamine™ 293 reagent were diluted with Opti-MEM™ I reduced serum media and incubated at room temperature for 5 min. After 15-20 min, ExpiFectamine™ 293/Plasmid DNA complexes were slowly added to Expi293F™ cells. Eighteen hours post-transfection, ExpiFectamine™ 293 Transfection Enhancer 1 and 2 were added to the transfected cells. Six days post-transfection, culture was harvested, and proteins were purified from culture supernatant by nickel affinity chromatography. The culture supernatant was incubated with Ni-Sepharose 6 Fast Flow resin (GE Healthcare) for 6-8 h at 4 °C. Non-specific proteins were removed by passing twenty-column volumes of wash buffer (PBS containing 25 mM Imidazole, pH 7.4). Bound proteins were eluted from the Ni-NTA column using 500 mM imidazole in PBS buffer, pH 7.4, and were dialyzed against PBS buffer using a 10 kDa (MWCO) dialysis membrane. The purity of purified protein samples was analyzed on SDS-PAGE.

### Size Exclusion Chromatography (SEC)

A Superdex-200 10/300 analytical column, equilibrated with PBS buffer, pH 7.4, was used for size exclusion chromatography-multi angle light scattering. 100-200 μg of purified protein samples were injected into the column, and protein peaks were resolved on a BioRad NGC chromatography system at a flow rate of 0.4 mL/min.

### Differential Scanning Fluorimetry (nano-DSF)

A Prometheus NT.48 instrument was used to determine the thermal stability of immunogens. The thermal unfolding of immunogens was monitored from 20 °C to 95 °C at a scan rate of 1 °C/min_59._

### Trypsin Proteolysis

The proteolytic stability of designed immunogens was studied by performing trypsin proteolysis. S2, RS2, and S2R were dialyzed against 50 mM Tris buffer (pH 7.5) containing 1 mM CaCl_2_, followed by incubation with TPCK trypsin (protease) in a 1:50 molar ratio at 4 °C and 37 °C. At different time points, aliquots were taken out, and the reaction was quenched using 6X-SDS-loading dye. Collected samples were analyzed on 12 % SDS-PAGE.

### Surface Plasmon Resonance (SPR)

The binding affinity of the S2, RS2, S2R and Spike immunogens to different RBD and S2-specific antibodies were measured using ProteOn XPR36 Protein Interaction Array V.3.1. The SPR sensor prism HC200M chip (Xantec bioanalytics) was first activated with EDC and sulfo-NHS. Then Protein G (10 μg/mL) was immobilized in the presence of 10 mM sodium acetate buffer, pH 4.0. Finally, excess sulfo-NHS esters were quenched using 1 M ethanolamine. 500 response units (R.U.) of ACE2-hFc, RBD specific monoclonal antibodies: CR3022, S309, ADG-2, H014, and S2-specific monoclonal antibodies: B6 and CC40.8 were immobilized. Different protein sample concentrations (100 nM, 50 nM, 25 nM, 12.5 nM, and 6.25 nM) were passed over the HC200M chip surface at a flow rate of 30 μL/min, followed by dissociation with PBS buffer containing 0.05% tween-20 (PBST). After each kinetic-binding assay, the chip was regenerated using 0.1 M Glycine-HCl, pH 2.7. Proteon Manager was used for fitting data to a simple 1:1 Langmuir interaction model to obtain kinetic parameters.

### Formulation preparation with SWE adjuvant

The adjuvant SWE (squalene-in-water emulsion) was co-developed by the Vaccine Formulation Institute (Switzerland) and Seppic (France) and is available at GMP grade (Sepivac SWE™) under an open access model. Immunogens (S2, S2R or RS2) were formulated with SWE at 1:1 volume ratio. The formulation RS2 in combination with SWE was evaluated for 1 month stability measuring adjuvant physicochemical characteristics and antigen integrity. Measurements included: visual inspection, particle size and polydispersity (DLS), zeta potential (ELS), pH, osmolality, and squalene content (HPLC). Antigen integrity was evaluated by ELISA.

### Immunization Studies

#### (BALB/c mice immunizations)

Groups of 6-8 weeks old, female BALB/c mice (n=5) were intramuscularly immunized with either 2 μg or 20 μg dose of SWE adjuvant formulated RS2 immunogen. *(hACE-2 expressing C57BL/6 transgenic mice immunizations):* Groups of hACE-2 expressing transgenic mice (n=6/7) were intramuscularly immunized with 2 μg and/or 20 μg dose of SWE adjuvant formulated immunogens (S2, RS2, S2R, Spike, and RBD). B6N; DBA2-Tg(K18-hACE2)3068Mgef/Blisc (acrc:21000509) mice strain was generated and provided by the Mouse Genome Engineering Facility, NCBS Bangalore using the K18-hACE2 transgene plasmid, kindly donated by Dr Paul B. McCray ^60^

#### (Syrian Hamster Immunizations)

Groups of five female golden Syrian hamsters were intramuscularly immunized with either a 5 μg or 20 μg dose of SWE adjuvant-formulated immunogens (RS2, Spike, and RBD). SWE adjuvant-treated animals were used as control. The sera samples from immunized and unimmunized animals were collected before prime (Day-1), 2 weeks post-prime (Day 14), and 14 days post-boost (Day 35) for endpoint ELISA and measuring neutralizing antibody titers. *Challenge Studies:* Twenty-one days following boost immunizations, immunized animals were intranasally challenged with either 10^5^ pfu MA10, 10^5^ pfu/ 10^4^ pfu Beta, or 10^4^ pfu Delta VOCs. The unimmunized challenged animals were used as control. Weight change of immunized-challenged animals, unimmunized-unchallenged animals (Unimmunized), and unimmunized-virus challenged control groups were monitored and recorded for 5-9 days. Post-challenge, lungs were harvested for viral titer estimation and histopathological examination were obtained on day 6 except for the lyophilized RS2 study, where lung samples were obtained on day 10, as previously described ^21^. For lung tissue histopathology scoring, we developed a scientific method using Mitchison’s virulence scoring system with some modifications, considering the consolidation of lungs, severity of bronchial and alveolar inflammation, immune cell influx, and alveolar and perivascular edema ^21,61^. The histopathology scores were graded as 0-4 (4: Severe pathology; 3: Moderate pathology; 2: mild pathology; 1: Minor/minimum pathology; 0: No pathology).

### ELISA and Competition ELISA

As described previously, endpoint titers of serum-binding antibodies were determined using ELISA ^22^. Briefly, 96 well ELISA plates were coated with 4 μg/mL RBD (332-532 a.a) or Spike (1-1208 a.a) and incubated at 25 °C for 2-3 h. Plates were washed with PBST, followed by incubation and with blocking with 3 % skimmed milk (in PBST) at 25 °C for 1 h. Four-fold serially diluted antisera raised against immunogens were added to wells and incubated at 25 °C for 1 h. Following three washes with PBST, plates were incubated with 1:5000 diluted goat ALP-conjugated anti-mouse IgG secondary antibody at 25 °C for 1 h. Plates were washed thrice with PBST and incubated with pNPP liquid substrate at 37 °C for 30 min. Optical density (O.D.) was measured at 405 nm. The ELISA endpoint titers were determined as the highest sera dilution with an O.D. signal above 0.2 at 405 nm. Competition ELISA was performed as previously describeds^22^. Briefly, 96 well plates (HIMEDIA, Cat# EP2-5X10NO) were coated with Expi293 cell produced RS2 at 4 µg/mL concentration in 1x PBS (60 µl/well) and incubated for 2 h at 25 °C under gentle shaking condition (300 rpm) on a thermomixer (Eppendorf, USA) and then plate was transferred to 4 °C cold room for overnight. Next day each well was washed with of 1xPBST (200µl/well) and then treated with blocking solution (100 µL 3% skimmed milk in 1xPBST) for 45 min at 25 °C, 300 rpm. The sera isolated from 5 animals in a group were used. Individual sera against the mentioned antigens were added at 2-fold serial dilution with a starting dilution of 1:25 in blocking solution (60µL). Only blocking solution was added to the control wells. The plates were then incubated for 1 h at 25°C, 300 rpm. Plates were provided with 3 additional washes with 1xPBST (200 µL of 1xPBST/well). An additional blocking step was also performed for 45 min with blocking solution (100µL) incubated at 25°C, 300rpm. An excess of monoclonal antibody either S2X259, or B6 were added (60µL at 20µg/mL) to their respective wells and incubated for one hour at 25°C, 300rpm. Next, three washes were given (200 µL of PBST/well) to remove excess unbound proteins. 50 µl/well Goat Anti-Human IgG Antibody, Alkaline Phosphatase conjugate (Sigma-Aldrich Cat # AP112A, Lot # 3519874; diluted 1:5000 in blocking buffer) was added and samples incubated for 1 hour at 25°C, 300 rpm. Plates were washed 3 times with 200 µL of PBST/well. Finally, 50 µL/well of a 37 °C prewarmed alkaline phosphatase yellow (pNPP) liquid substrate (Sigma-Aldrich, Cat # P7998, Lot # SLCJ1764) was added, and plates were incubated for 30 minutes at 37 °C, 300 rpm. The chromogenic signal was measured at 405 nm using an ELISA plate reader (Emax-Plus Microplate reader, Molecular Devices).

The percent competition was calculated using the following equation.

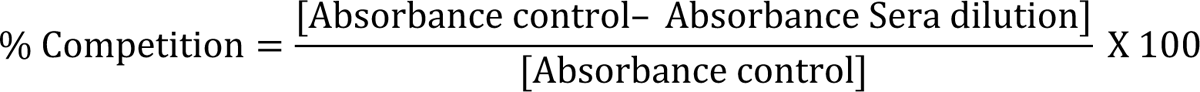

where, *Absorbance control* is the absorbance at 405nm of ACE2-hFc, S2X259 or B6 protein/Antibody binding directly to RS2 protein in the absence of sera, Absorbance *sera dilution* is the absorbance where the serum dilution is incubated with competitor agents like S2X259 or B6 protein. The % competition as a function of serum dilution was fit using a three-parameter nonlinear least squares fit curve using Graph Pad Prism 10.0. The sera dilution at 50 % competition on the fitted curve was termed as the IC_50_ competition titer.

### Pseudoviral neutralization assay

Psuedoviral neutralization assays were performed as previously described ^21^. The genes encoding Spike proteins from VOC were synthesized at GenScript (USA) except for the SARS-CoV-1 pseudovirus which was obtained from Dr Kalpana Luthra at the All India Institute of Medical Sciences, New Delhi. Pseudovirus neutralization titers (ID_50_) were defined as the serum dilution at which the infectivity of the virus is reduced to 50%.

### Statistical Analysis

Data analysis was performed using GraphPad Prism software 10.0.0. The ELISA binding, neutralization titers, and pseudoviral virus neutralization titer data were analyzed with a two-tailed Mann–Whitney test and non-parametric Kruskal–Wallis with Dunn’s multiple, respectively. Weight changes in mice and hamsters were analyzed with a two-tailed Student’s t-test. (* indicates *p* < 0.05, ** indicates *p* < 0.01, *** indicates *p* < 0.001, **** indicates *p* < 0.0001). All mice and Hamster immunization studies were approved by the Institutional Animal Ethics Committee (CAF/ETHICS/847/2021; CAF/ETHICS/887/2022). These were carried out at the Central Animal Facility (CAF), Indian Institute of Science, according to CPCSEA and ARRIVE guidelines.

## Acknowledgments

NM acknowledges the Prime Minister Research Fellowship for her fellowship (PM/MHRD-20-17303.03). RV is a JC Bose Fellow of DST. DC acknowledges the Council of Scientific and Industrial Research for his fellowship (09/079(2837)/2019-EMR-I). We acknowledge Ms. Simran Srivastava for providing the sarbecoviruses spike constructs.

## Author contributions

Conceptualization: RV, NM

Methodology: RV, NM, SK, RSR, RS, SBJ, MB, DC, SP, RPR, CL, VJ, PMD, AJ, SSSA

Investigation: NM, SK, RSR, RS, SBJ, NJ, MB, DC, SP, RPR, CL, VJ, Visualization: NM, SK, RSR, RS, SBJ, NJ, MB, DC, SP, RPR, CL, VJ

Funding acquisition: RV Project administration: RV Supervision: RV

Writing – original draft: NM

Writing – review & editing: All authors

## Data and materials availability

All data are available in the main text or the supplementary materials.

## Supplementary Figure Legends

**Fig. S1.**
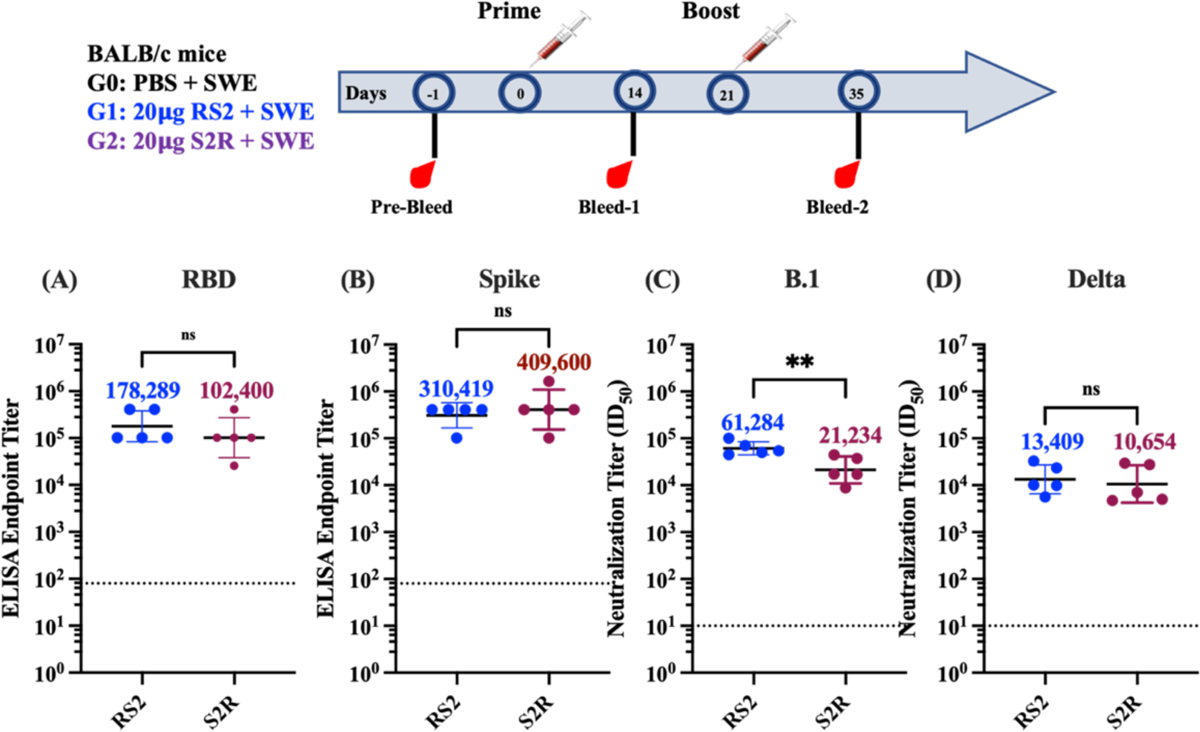
Comparative immunogenicity of RS2 and S2R in mice. BALB/c were mice immunized twice with 20µg of RS2 or S2R. Two weeks following boost vaccination RBD and Spike specific IgG levels and neutralizing titers were measured in immunized mice sera samples. Comparative ELISA endpoint titers against **(A)** RBD and **(B)** Spike ectodomain, respectively **(C and D)** Comparative neutralizing antibody titers elicited by 20µg of RS2 and S2R against **(C)** B.1, **(D)** Delta variant pseudoviruses. Titers are shown as geometric mean with geometric SD. The ELISA binding and neutralization titers were analyzed with a two-tailed Mann–Whitney test. (ns indicates non-significant, * indicates *p* < 0.05, ** indicates *p* < 0.01, **** indicates *p* < 0.0001).

**Fig. S2.**
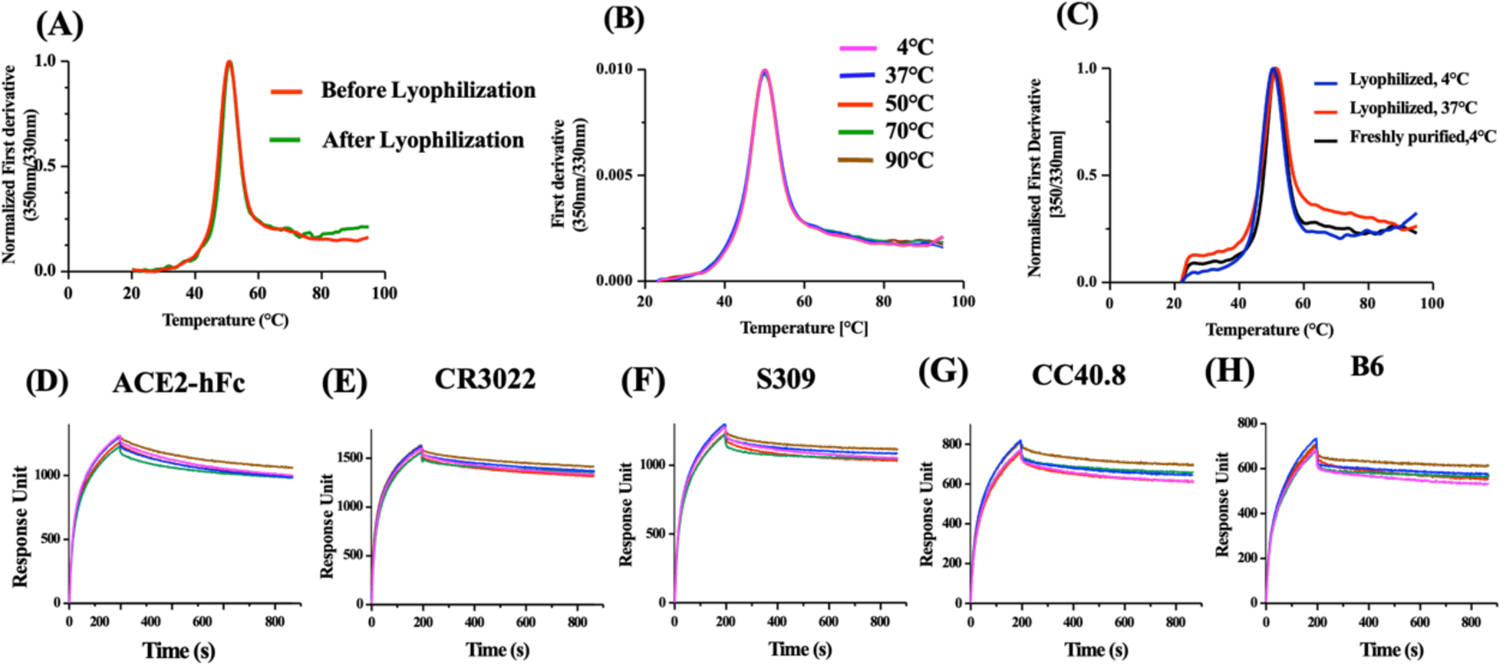
Effect of transient thermal stress on lyophilized RS2. RS2 protein was dialyzed in water and then lyophilized. **(A)** Equilibrium thermal unfolding profile of RS2 before and after lyophilization and resolubilization in PBS. Lyophilized RS2 was incubated for one hour at different temperatures (4, 37, 50, 70 and 90℃). Following reconstitution in PBS buffer, thermal stability of **(B)** RS2 was monitored using Nano-DSF. **(C)** Thermal stability of lyophilized and resolubilized RS2 after incubation at 37 ℃ for 1 month. **(D-H)** In addition, RS2 samples were characterized for binding of with panel of RBD **(D-F)** and S2 **(G and H)** specific antibodies using SPR **(D)** ACE2-hFc, **(E)** CR3022, **(F)** S309, **(G)** CC40.8, and **(H)** B6.

**Fig. S3.**
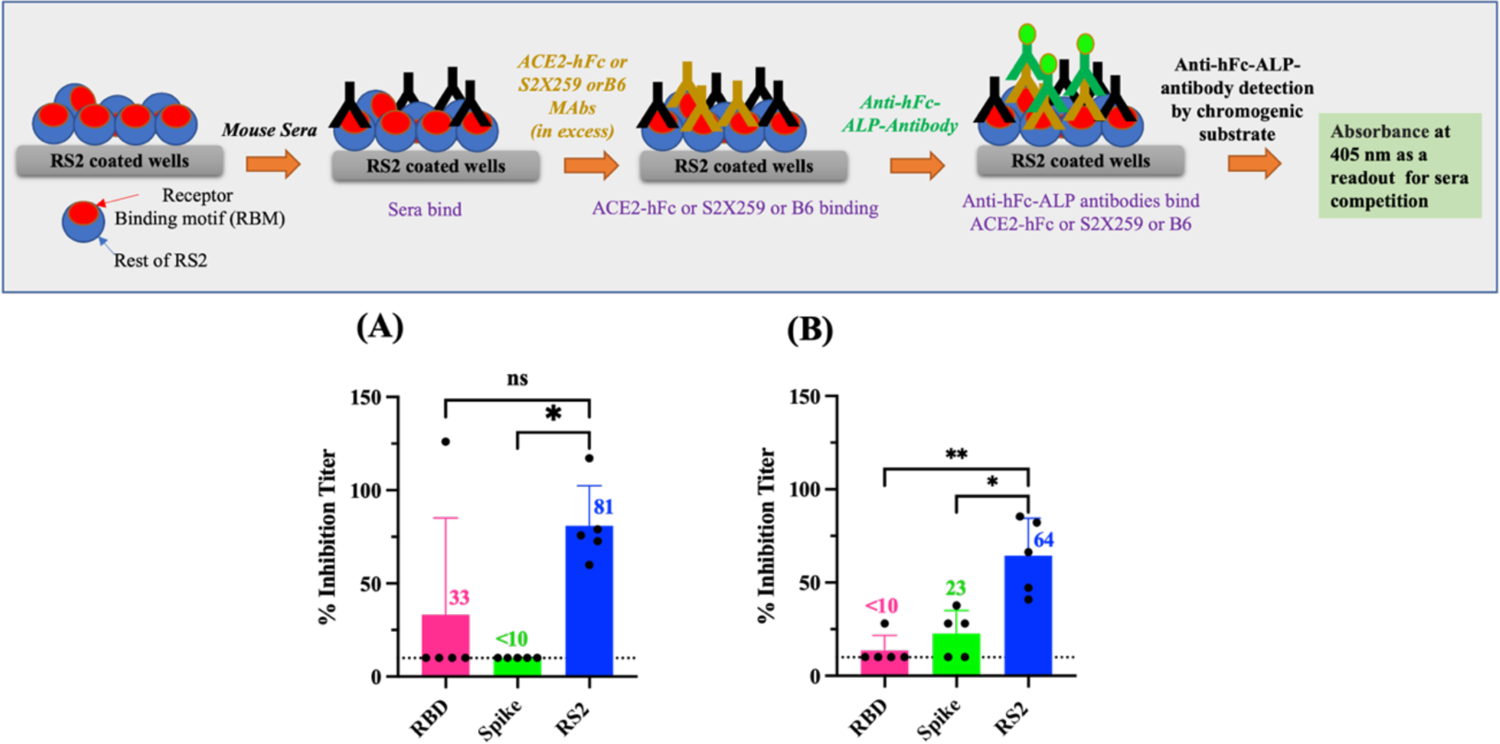
Competition of immunized mice sera with RBD and S2 specific monoclonal antibodies. BALB/c mice were immunized with 20µg of RBD, Spike, or RS2. Top panel: Schematic of assay. Bottom panel **(A and B)** ELISA competition titers against **(A)** S2X259 (Class 4) and **(B)** B6 (S2 helix) monoclonal antibodies for individual, immunized mice sera. Immunogens used to elicit respective sera are listed on the x-axis and competition titers on the y-axis of each bar graph.

